# System and transcript dynamics of cells infected with severe acute respiratory syndrome virus 2 (SARS-CoV-2)

**DOI:** 10.1101/2024.03.25.586528

**Authors:** João M. F. Silva, Jose Á. Oteo, Carlos P. Garay, Santiago F. Elena

## Abstract

Statistical laws arise in many complex systems and can be explored to gain insights into their structure and behavior. Here, we investigate the dynamics of cells infected with severe acute respiratory syndrome virus 2 (SARS-CoV-2) at the system and individual gene levels; and demonstrate that the statistical frameworks used here are robust in spite of the technical noise associated with single-cell RNA sequencing (scRNA-seq) data. A biphasic fit to Taylor’s power law was observed, and it is likely associated with the larger sampling noise inherent to the measure of less expressed genes. The type of the distribution of the system, as assessed by Taylor’s parameters, varies along the course of infection in a cell type-dependent manner, but also sampling noise had a significant influence on Taylor’s parameters. At the individual gene level, we found that genes that displayed signals of punctual rank stability and/or long-range dependence behavior, as measured by Hurst exponents, were associated with translation, cellular respiration, apoptosis, protein-folding, virus processes, and immune response.

**Author summary:** Viruses replicate within susceptible cells by exploiting the cellular machinery. Consequently, cells initiate defenses against the virus and signal other cells, notably immune cells. This ongoing battle prompts significant alterations in the cells’ gene expression patterns throughout the infection process. In this study, we apply statistical principles from complex systems theory to analyze gene expression data from individual cells infected with SARS-CoV-2. Our research aims to elucidate how viral infection impacts cells at both systemic and individual gene levels. Our primary findings are twofold: (*i*) the virus influences the distribution of gene transcripts over the course of infection, varying depending on cell type. (*ii*) As the infection progresses, numerous genes associated with critical cellular functions and immunity exhibit signs of punctual instability and/or autocorrelation, indicating their response to viral infection at various stages of the process.

## Introduction

Transcriptomics analyses commonly rely on linear models to test whether the mean expression of any set of genes is altered in response to a treatment or condition, which are usually treated as factors in the model. Although changes in gene expression level are of the utmost importance in Biology, aspects about the whole system behavior are not captured by these models. Another limitation of linear models is that by treating conditions as factors might lead to the loss of important information about the time course variation of transcripts. For instance, in single-cell RNA-sequencing (scRNA-seq) data, cells from the same cell type can be at distinct differentiation stages during sample preparation. Thus, the inference of a continuous pseudotime trajectory of the transition from one cell type/stage to another, where each cell is assigned a value based on its relative position along it, can provide a continuous covariate for statistical models with higher sensitivity than factors to identify differentially expressed genes [1,2]. Accordingly, it has been recently shown that along the progression of cellular infection, the response to severe acute respiratory syndrome virus 2 (SARS-CoV-2) is triphasic, and that treating infected cells as one factor in an infected *vs*. uninfected linear model will lead to biases in identifying differentially expressed genes [3].

Several statistical models and frameworks have been applied to transcriptomics data to model its structure and disentangle useful biological information from sampling noise and/or intrinsic stochastic biological variation. A recent study identified various emerging statistical laws from complex compartment systems on scRNA-seq data [4]. While the negative binomial distribution is often used to model both scRNA-seq and bulk RNA-seq count data, scRNA-seq data present some unique characteristics. The low capture rate of transcripts in scRNA-seq experiments makes that only about 10 - 20% of the transcripts from each cell are sequenced [5,6]. Due to this phenomenon, known as dropout, where a gene is expressed in a cell but its transcripts are not captured, gene count matrices from this type of experiments are sparse. Protocols for the preparation of scRNA-seq data also often rely on unique molecular identifier (UMI) tags that are added to the transcripts during RT-PCR to drastically reduce amplification bias [5,7–9].

The dynamics of various cellular processes can be explored with statistical models from complex systems. For instance, power law relationships arise naturally in many complex systems, including in scRNA-seq data [4]. In particular, an empirical law known as Taylor’s law states that there is a power relationship between the mean of an element *x* and its standard deviation in the form of *σ* = *V*⟨*x*⟩^β^ [10]. If *β* = 0.5, then the system dynamics follows a Poisson distribution, and if *β* = 1, then the system fits to an exponential distribution, meaning that its elements are aggregated [10–12]. In the special case of time series data, Taylor’s parameter *V* has been used as a proxy of system stability through time for data from the human microbiome [12]. In log-log scale, *V* is the intercept term. Whenever *V* is large, the standard deviation of each element in the system will also be large, a fact that is associated with system instability.

The long-range dependence of a time series is a feature that has been thoroughly studied with the so-called Hurst’s rescaled range analysis [13–15]. Records in time are associated to an index *H*, known as Hurst exponent, that runs between zero and one and, importantly, has an interesting interpretation. Values of *H* > 0.5 convey that the temporal sequence presents persistence. This is a kind of bias which means that the future variations tend to be similar to the past ones in the sequence. Antipersistence (*H* < 0.5) is defined the other way around. Hurst found empirically that a large number of natural processes studied with the rescaled range yield *H* values close to 0.7, which is termed *Hurst phenomenon* in the literature. This analysis has been applied to temporal transcriptomic data from *Escherichia coli* and *Saccharomyces cerevisiae*, where it was shown that most genes exhibited *H* > 0.5 values, which are indicative of persistent long-range dependence [16]. Also, in a recent study, the rescaled range analysis shows the persistent character of the distribution of mutations along human chromosomes [17].

Here, we aim to gain insights into the structure and system behavior of cells infected with SARS-CoV-2 along the course of infection. We first hypothesize that the rank dynamics of transcripts and system behavior of scRNA-seq data from infected cells can be explored by fitting gene abundances to Taylor’s law. We find that, in all cases, fluctuations grow with mean value on a biphasic Taylor’s law, consisting in a Poisson and an exponential laws separated by a breaking point. Both, progression of infection and sampling noise have a significant impact on the estimation of Taylor’s parameters. The rank dynamics of a gene gives us information about its relative importance in the system. For each gene, we investigate whether its rank is stable and calculate their associated Hurst exponent along the course of infection. The robustness of these methods was further assessed by the use of control datasets. Overall, we found evidence of several genes exhibiting punctual rank stability and/or persistent behavior that are related to viral processes or immune responses that could serve as potential pharmaceutical targets for the treatment of COVID-19.

## Results

### Detection of SARS-CoV-2 genome and identification of infected cells

The presence of viral RNA was investigated in four datasets from human bronchial epithelial cells (hBECs) [18] and six human intestinal epithelial cells (hIECs) [19], divided in three datasets from colon and three from ileum organoids. Viral RNA was detected in all datasets, although the detection of SARS-CoV-2 RNA in 28 mock-infected hBECs, 11 colon cells and 4 ileum cells are likely due to misalignments. To differentiate between infected cells supporting viral activity from droplets that contained viral RNA from attached viral particles or ambient viral particles or RNA, we sought to estimate the mean SARS-CoV-2 UMI count from empty droplets to set a threshold for calling infected cells. However, no viral RNA was detected in the empty droplets, and thus, a threshold of 10 viral UMIs was set. With this strategy, 1%, 8.5% and 11.5% of hBECs were infected at 1, 2 and 3 dpi, respectively; 11.5% and 96.3% colon cells were infected at 12 and 24 hpi, respectively; and 23.9% and 95% ileum cells were infected at 12 and 24 hpi, respectively. A high proportion of infected cells was observed, in particular for hIECs. This is in contrast with the number of infected cells in these datasets reported in [19], where the infection rate was estimated to be lower than 10%. Despite this, we decided to follow with our strategy due to the following two reasons. First, in the original work with the hIECs datasets, the proportion of infected cells was estimated based on immunofluorescence staining of dsRNA and SARS-CoV-2 N protein [19]. It is likely that, at the beginning of infection, viral replication is low. Therefore, dsRNA might not be readily detected and N protein translation might also be low or even not yet synthesized. This means that many infected cells at the beginning of infection might be missed by immunofluorescence staining. Second, keeping uninfected cells in our analyses should not compromise our results. Instead, it would just make that the infection trajectory starts with uninfected cells, irrelevant for the application itself of statistical laws from complex systems to pseudotemporal scRNA-seq data.

### Transcript abundances fit a biphasic Taylor’s law

Gene abundances data of SARS-CoV-2-infected cells are represented in the Taylor plot of Fig 1. The distribution of points suggests a fit to a biphasic, or segmented, linear regression whose outcomes are in Table 1. The data from all three cell types fit better to the biphasic model than to an unsegmented model with no breakpoint (*F*-tests in Table 1). In the biphasic model, two *V* and two *β* parameters are estimated, where *V*_1_ and *β*_1_ are the ones estimated for the data points below the breakpoint and *V*_2_ and *β*_2_ the ones estimated for the data points above the breakpoint. For larger abundances, we find a slope *β* ≈ 1, characteristic of the exponential distribution. For smaller abundances, we find a slope *β* ≈ 0.5, characteristic of the Poisson distribution. This biphasic behavior in the Taylor plot is most likely related to sampling noise for the low capture rate of transcripts in scRNA-seq protocols [20]. Additionally, the rank stability index (*RSI*; see *Methods*) was calculated for each gene. Higher rank stability seems to be associated to high expressed genes that follow an exponential distribution (Fig 1).

**Fig 1.**
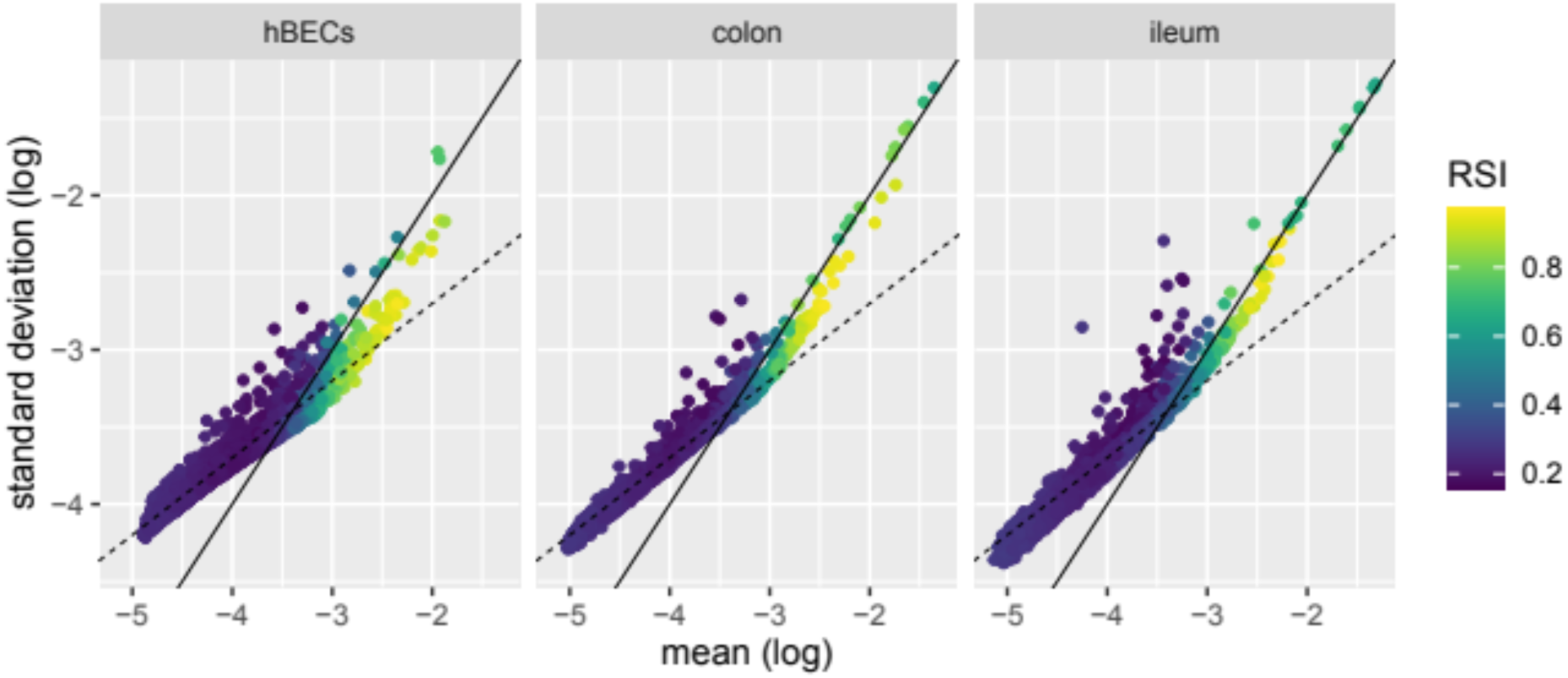
Taylor’s law plots. For illustrative purposes, solid lines correspond to the exponential distribution (*β* = 1 and log(*V*) = 0), and dashed lines to the Poisson distribution (*β* = 0.5 and log(*V*) = –1.7), where log(*V*) was chosen to be –1.7 for better visualization. The *RSI* of each gene is shown.

**Table 1.** Parameters of the segmented fit to Taylor’s law for infected cells. Matrix sparsity (proportion of zeros), number of genes and number of cells are also shown.

|  |  | hBECs | colon | ileum |
| --- | --- | --- | --- | --- |
| Matrix properties | sparsity | 0.77 | 0.76 | 0.72 |
|  | Number of genes | 7978 | 7567 | 8333 |
|  | Number of cells | 3207 | 4703 | 4433 |
| $F$-test<sup>1</sup> | $F$ | 0.82 | 0.53 | 0.76 |
| | $P$ | < 0.0001 | < 0.0001 | < 0.0001 |
| Segmented fit | $V_1$ | 0.02 | 0.03 | 0.04 |
| | $\beta_1$ | 0.48 | 0.53 | 0.57 |
| | $V_2$ | 0.25 | 1.47 | 1.28 |
| | $\beta_2$ | 0.9 | 1.12 | 1.07 |
|  | breakpoint (log(mean)) | -3.16 | -2.99 | -3.07 |

<sup>1</sup>One-tailed $F$ -tests of the residuals of the segmented vs. unsegmented fits

### The simulated control datasets mimic the increase in sampling noise seen in infected cells

In order to conduct a more thorough analysis of the progression of infection, infected cells were divided into bins with cells showing a similar viral load. We found that 30 bins were a good compromise between number of bins and number of cells in each bin. As infection progresses and cells accumulate more viral RNA, the sampling of cellular transcripts become compromised and a higher incidence of zeros in the matrix due to dropout is seen. Therefore, to better understand the effect of dropout in the identification of temporal signal embedded in scRNA-seq data, we simulated the expected increase in sampling noise as infection progresses by down-sampling UMIs from uninfected cells. The proportion of zeros per gene per bin of the simulated control datasets follows the same trend as true infected cells (S1A Fig). To investigate whether the simulated dataset retains the same transcriptional profile from the uninfected cells, we performed standard clustering analysis with cells from the simulated dataset and their “matching” uninfected cell. Cells from the simulated datasets clustered together with uninfected cells confirming that they still retain the same transcriptional profile (S1B Fig).

### Analysis of infection progression reveals signals of varying system dynamics

To further investigate system stability throughout infection, Taylor’s parameters were estimated for each bin (see section above) in the three cell types. An increase-decrease-increase pattern in both *V* and *β* was observed in hIECs (Fig 2A). In these cells, *V* and *β* increase with sampling noise for the simulated dataset, in contrast to the pattern seen in infected cells (Fig 2A). On the contrary, a similar pattern between the infected and simulated datasets was seen for hBECs, with an initial oscillation and a sharp increase of parameters *V* and *β* at the end of the infection.

**Fig 2.**
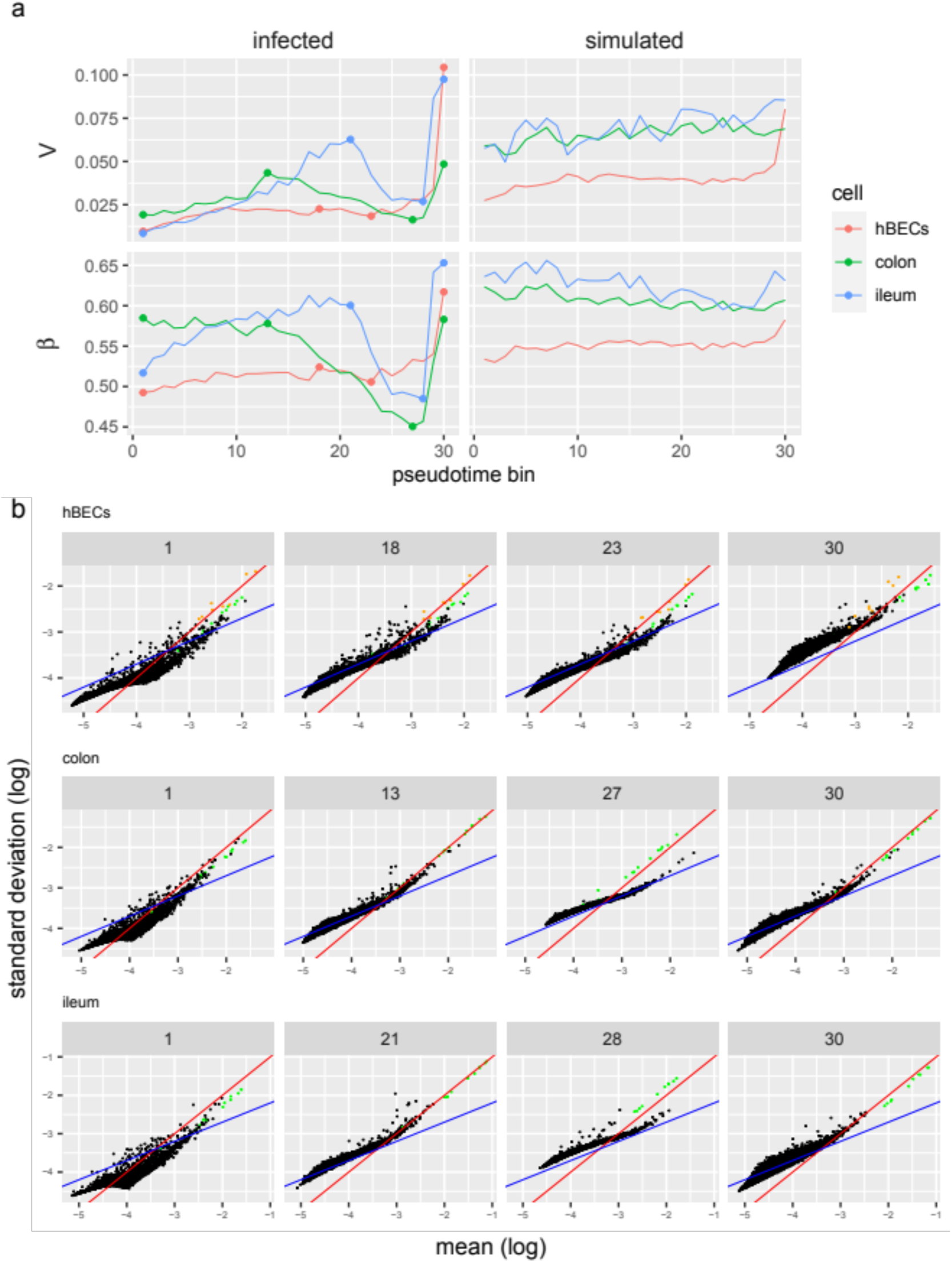
Evolution of Taylor’s parameters along the infection. (a) Taylor’s parameters estimated from each bin for infected cells and the simulated datasets. Dots represent selected bins which fits to Taylor’s law are shown in (b), where mitochondrial genes are shown in green; and for hBECs, selected nuclear genes that generally followed an exponential distribution regardless of rank are shown in orange. Red lines correspond to the exponential distribution (*β* = 1 and log(*V*) = 0), and blue lines to the Poisson distribution (*β* = 0.5 and log(*V*) = –1.7), where log(*V*) was chosen to be –1.7 for better visualization.

Mitochondrial expressed genes show an exponential distribution, even for lower abundances (Fig 2B; bins 27 and 28 for colon and ileum cells, respectively). This suggests that they are not responding to infection, but rather their rank is shifting due to differences in the expression of other genes. Detailed inspection of the plots shows that, for hBECs, some nuclear genes, most notably *LCN2*, *S100A2*, *S100A9*, *SCGB1A1*, *SCGB3A1*, *SERPINB3*, *SLPI*, and *WFDC2*, generally follow an exponential distribution regardless of their rank, suggesting that, like mitochondrial genes, they are aggregating in most bins and may not be responding to infection. We can observe how the thickness in the distribution of points in the Taylor’s plot changes with the infection process (Fig 2B). This effect is due to the sparsity of the gene counts, which grows with infection (S1A Fig). If the gene counts matrix contains an even number of zero and nonzero counts, a bell shape distribution of bins is observed (beginning of infection). Otherwise, if the gene counts matrix contains a dominant number of zero counts, the shape distribution of bins is much thinner.

Next, we performed a segmented fit to Taylor’s law for each bin to estimate Taylor’s parameters in the biphasic regime. Biphasic Taylor’s parameters *V*_1_ and *β*_1_, that fit to gene abundances with a Poisson behavior, exhibited a similar pattern to the unsegmented fit parameters *V* and *β* (S2 Fig). Notably, *β*_1_ was lower than *β* at the beginning of infection, although *V*_1_ and *β*_1_ exhibited the same increase-decrease-increase behavior of *V* and *β* for hIECs. As infection progresses, the breakpoint increases for all three cell types (S2 Fig).

To further ascertain that the observed changes in Taylor’s parameters are not due to technical noise, we performed ANCOVA tests for the effect of infection progression (herein pseudotime; where each bin corresponds to a different point throughout the infection), cell type and their interaction on each Taylor parameter, while also adding the number of genes and matrix sparsity as covariates to control for increasing sampling noise in the system. The rationale of adding these covariates is that as less cellular transcripts are captured due to increasing viral RNA accumulation a higher proportion of zeros will be observed and less genes will have their transcripts captured. All explanatory variables, with the exception of sparsity for *β*, had a significant effect on parameters *V* and *β*. The largest effect sizes (partial *η*^2^) were estimated for cell type and the interaction between pseudotime and cell type for parameter *V* and cell type, number of genes and the interaction between pseudotime and cell type for *β* (Table 2). Additionally, we performed these analyses on the simulated control datasets. Whereas sampling noise had a significant effect on Taylor’s parameters for the simulated datasets, its interaction with cell type was not significant for *V* and, although significant for parameter *β*, its effect size was lower than that of the infected dataset (Table 2). From these analyses we conclude that noise induced by dropout has a uniform effect on Taylor’s parameter *V* and a cell type-dependent effect on parameter *β*, and that infection with SARS-CoV-2 induces changes in the distribution properties of the system that is also dependent on cell type.

**Table 2.** *P*-values from ANCOVA analysis of Taylor’s parameters for cells infected with SARS-CoV-2 and the simulated dataset. Partial *η*^2^ values are shown in parenthesis (partial *η*^2^ ≥ 0.15 are conventionally taken as large effects).

| | $V$ | | $\beta$ | |
| --- | --- | --- | --- | --- |
| Effect | infected | simulated | infected | simulated |
| Pseudotime | < 0.0001 (0.33) | < 0.0001 (0.44) | 0.0096 (0.08) | < 0.0001 (0.19) |
| Cell | 0.0003 (0.18) | < 0.0001 (0.85) | < 0.0001 (0.46) | < 0.0001 (0.94) |
| Sparsity | 0.0006 (0.13) | 0.0187 (0.07) | 0.87962 ($2.81 \times 10^{-4}$) | < 0.0001 (0.46) |
| Number of genes | 0.0016 (0.11) | 0.8539 ($4.16 \times 10^{-4}$) | < 0.0001 (0.52) | 0.0319 (0.05) |
| Pseudotime-by-cell | 0.0002 (0.19) | 0.1033 (0.05) | < 0.0001 (0.45) | 0.0014 (0.15) |

### Genes that display punctual rank stability are, most notably, related to translation, protein folding, and apoptosis

Genes exhibiting punctual rank stability were found by evaluating whether its *RSI* value (calculated from its mean expression at each bin; see *Methods*) is higher than expected by chance, irrespective of whether its *RSI* was high or low. The *RSI* and punctual stability index (*PSI*; see *Methods*) of each gene is shown in Fig 3A. In total, 380, 2840 and 4230 genes were found to exhibit punctual rank stability in hBECs, colon and ileum cells, respectively (S1 File). To ascertain that these results are robust, we also applied this approach to the simulated control datasets and uninfected cells, where four, eight and six false-positives were detected in hBECs, colon and ileum cells simulated datasets, respectively; and 15, 10 and 11 false-positives were detected in uninfected hBECs, colon and ileum cells, respectively (S1 File). The low false-positive rate of this analysis indicates our results are robust, and that the higher number of genes displaying punctual stability in hIECs might be related to intrinsic differences in the response to viral infection between hBECs and hIECs.

**Fig 3.**
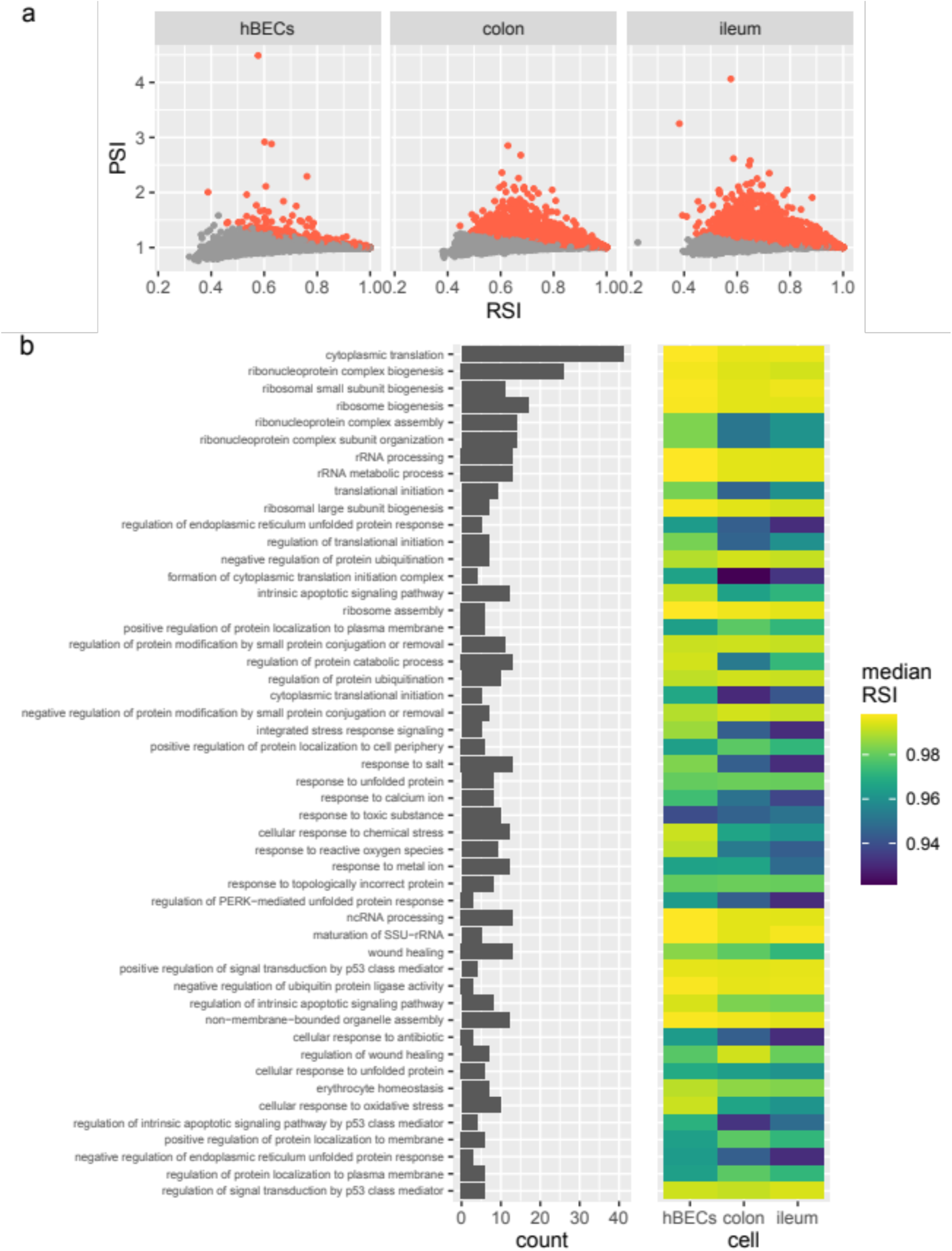
Gene rank stability dynamics. (a) Comparison of *RSI* with *PSI* of all genes for hBECs, colon and ileum cells. Each point represents one gene. Genes with significant punctual stability (FDR < 0.05) are shown in red. (b) Top 50 enriched GO terms, ranked by *P*-value, for genes that displayed signal of punctual rank stability in all three cell types. The median *RSI* of the genes in each GO term is shown for each cell type.

Punctual rank stability in all three cell types was found for 172 genes. GO terms enrichment analysis of these genes revealed an enrichment in those related to cytoplasmic translation (GO:0002181), regulation of apoptotic signaling pathway (GO:2001233), regulation of endoplasmic reticulum unfolded protein response (GO:1900101), and a few terms related to innate immunity such as response to lipopolysaccharide (GO:0032496), among others (Fig 3B; S1 File). Those genes that showed signal of punctual rank stability in all three cell types were generally stable, as shown by their median *RSI* (Fig 3B; S1 File). Among these, genes associated with translational processes and gene expression, such as translational elongation (GO:0006414), ribosome assembly (GO:0042255), maturation of SSU-rRNA (GO:0030490) and ncRNA processing (GO:0034470) were the most stable; and those associated with protein-DNA complex subunit organization (GO:0071824), detoxification (GO:0098754), epithelial cell apoptotic process (GO:1904019) and processes associated with immune response, such as response to lipopolysaccharide (GO:0032496), response to molecule of bacterial origin (GO:0002237) and myeloid leukocyte migration (GO:0097529) were the least stable (Fig 3B; S1 File).

### Several genes related to translation, cellular respiration, and viral processes, show evidence of persistent rank behavior

Next, we examined the presence of persistent behavior of gene rank along the course of infection, which indicates whether a gene has a tendency to maintain its rank once it changes. The robustness of the estimation of *H* from rank data was assessed by performing the analyses on a set of control datasets that included the simulated data, random matrices, uninfected cells, and shuffled infected cells that are not ordered according to viral RNA accumulation. The Hurst exponents calculated from these control datasets seemed to follow a normal distribution with a mean close to ∼0.5 which is expected for data with no temporal correlation, with the exception of the simulated ileum dataset that showed a small deviation towards higher *H* values (Fig 4A). Infected cells ordered according to viral RNA accumulation displayed a broad distribution of *H* values whose mean were nonetheless visibly higher than those from the control datasets (Fig 4A). By analyzing *H* values inside the Taylor’s law plot for each dataset of infected cells, we found that low expressed genes tended to exhibit slightly higher *H* exponents, most noticeably for ileum cells (Fig 4B). Given that gene rank was randomized in the case of ties, and that low expressed genes will be tied when their expression is zero, the higher *H* values of these low expressed genes is most likely artificially inflated.

**Fig 4.**
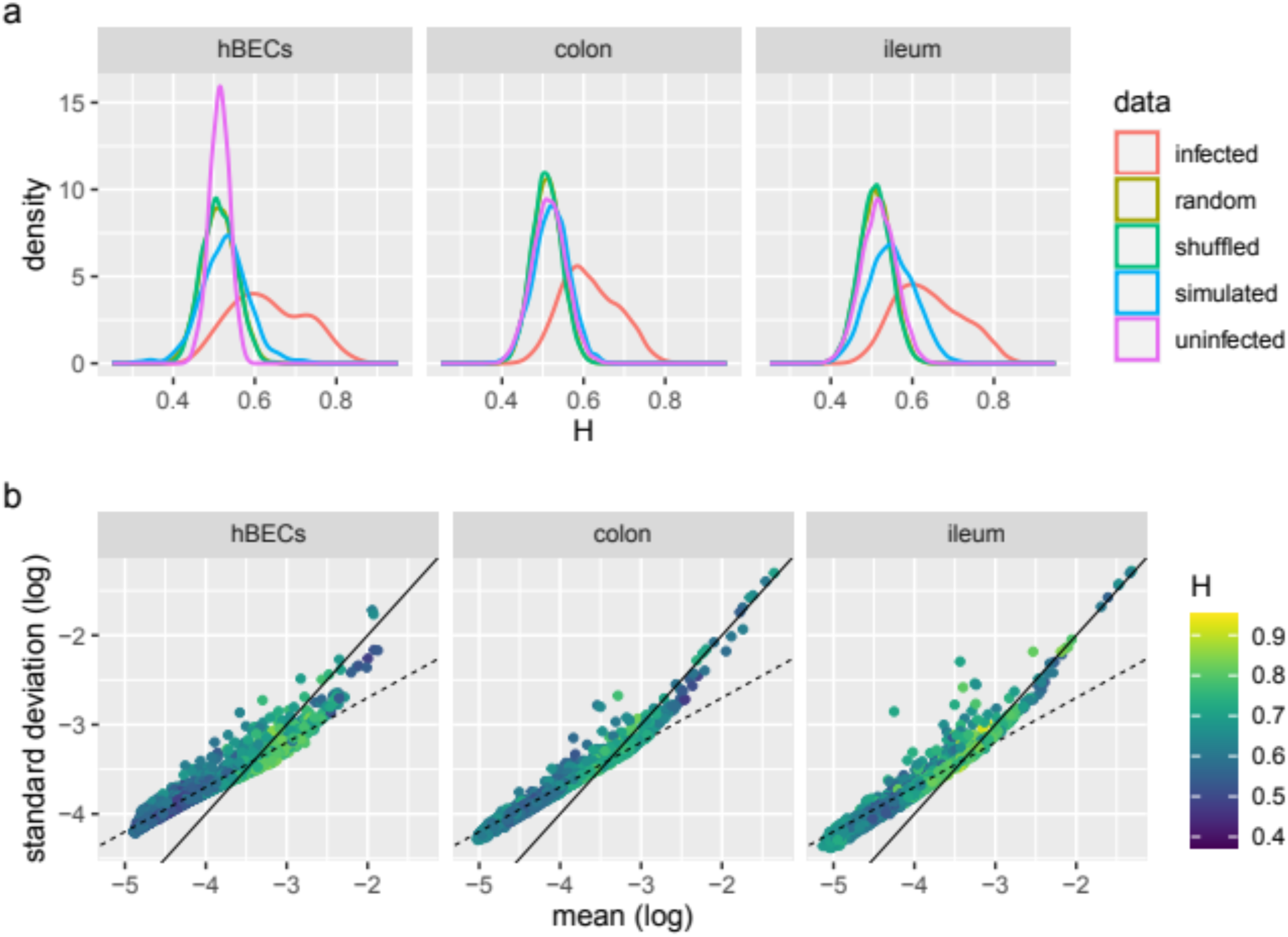
Estimation of H exponents from rank data. (a) Kernel density estimation of *H* exponents estimated from gene rank data from infected cells and control datasets for hBECs, colon and ileum cells. (b) Taylor’s law plots showing the *H* value of each gene for each cell type. Solid lines correspond to the exponential distribution (*β* = 1 and log(*V*) = 0), and dashed lines to the Poisson distribution (*β* = 0.5 and log(*V*) = –1.7, chosen for better visualization).

Based on the observations above, genes exhibiting strong persistent rank behavior were determined by comparing its *H* exponent from infected cells *vs*. the one calculated from the simulated dataset. Overall, the *H* exponents from infected cells tended to present higher values than their corresponding exponents from the simulated dataset (Fig 5A). We observed that 610, 657 and 1569 genes, out of 9910, 7582 and 8333, showed evidence of strong persistent rank behavior (*H* ≥ 0.7 in infected cells and, concomitantly, *H* < 0.7 in simulated dataset) in hBECs, colon and ileum cells, respectively. As discussed above, a higher false-positive rate in ileum cells is expected given that low expressed genes showed slightly higher *H* values.

**Fig 5.**
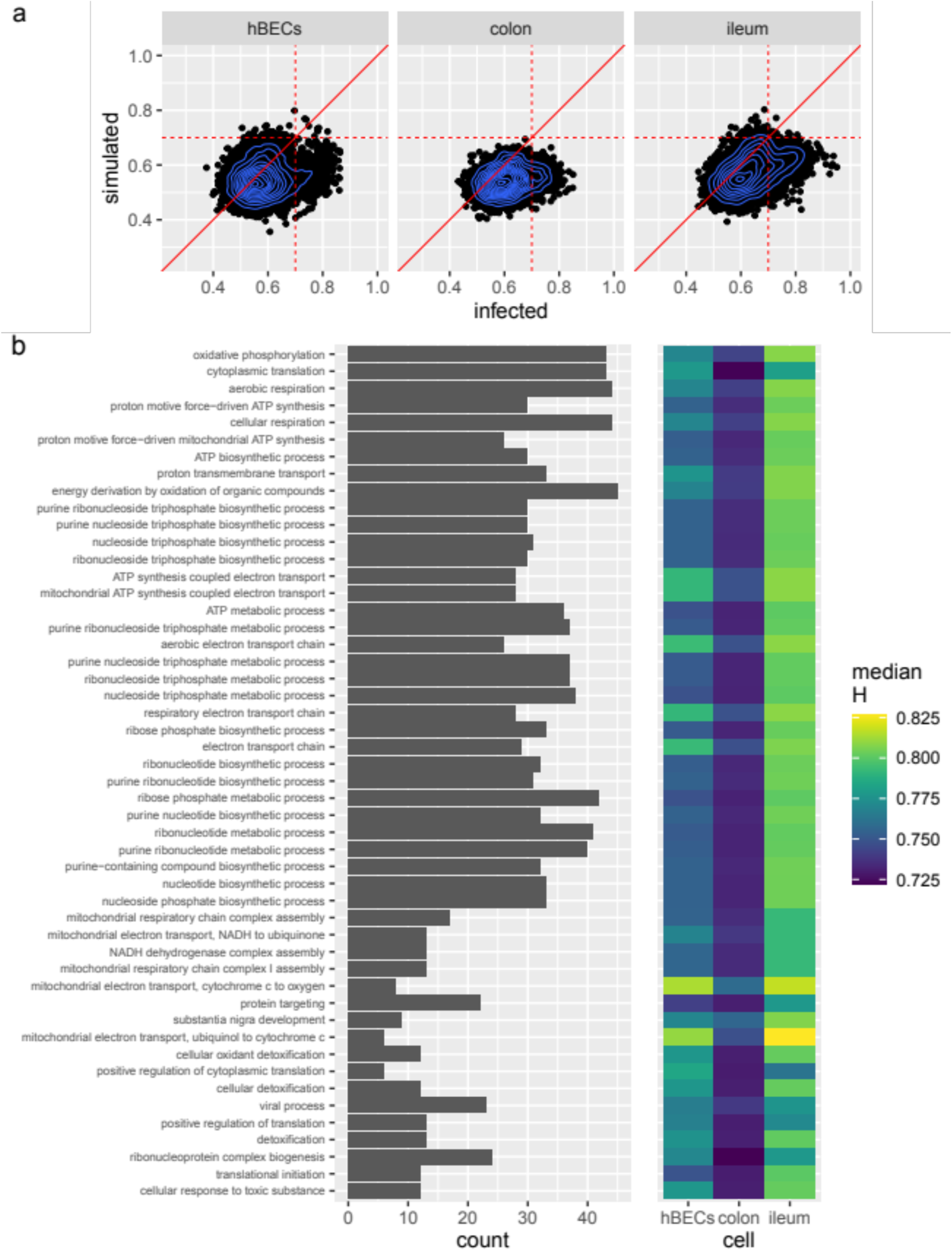
Functional analysis of genes exhibiting strong persistent behavior. (a) Comparison of the *H* exponents from infected and simulated data for hBECs, colon and ileum cells. Each point represents one gene. Dashed red lines show the *H* = 0.7 (Hurst phenomenon) threshold for evidence of strong persistent behavior, and the solid red lines are bisecting lines. Blue lines represent kernel density of the data. (b) Top 100 enriched GO terms, ranked by *P*-value, for genes exhibiting evidence of persistent rank behavior in all three cell types. The median *H* of the genes in each GO term is shown for each cell type.

In total, 297 genes displayed evidence of strong persistent rank behavior concomitantly in all three cell types. Amongst those, we found an enrichment of genes related to cytoplasmic translation (GO:0002181), cellular respiration (GO:0045333) and processes related to viral infection, such as internal ribosome entry site (IRES)-dependent viral translational initiation (GO:0075522), viral process (GO:0016032), viral translation (GO:0019081), and viral life cycle (GO:0019058), among others (Fig 5B; S2 File). An enrichment of genes related to IRES-dependent viral translation is unexpected since SARS-CoV-2 is not known to contain an IRES [21]. The median *H* of the genes in these GO categories varied little between 0.7 and ∼0.8, and were overall higher in ileum cells and lower in colon cells (Fig 5B; S2 File). In general, processes related to cellular respiration, such as mitochondrial electron transport, cytochrome c to oxygen (GO:0006123), mitochondrial electron transport, ubiquinol to cytochrome c (GO:0006122) and aerobic electron transport chain (GO:0019646) exhibited higher median *H* values, together with other processes such as mRNA stabilization (GO:0048255), regulation of substrate adhesion-dependent cell spreading (GO:1900024) and negative regulation of oxidative stress-induced intrinsic apoptotic signaling pathway (GO:1902176).

## Discussion

The underlying characteristics and dynamics of complex systems can be captured by several simple statistical laws. Here, we focus on a dynamical complex system of cells infected with SARS-CoV-2 to uncover how the system behaves as a function of infection progression. First, we fitted transcript abundance data to Taylor’s law to study system-level dynamics as previously done with data from the human gut microbiome [12]. A biphasic fit to Taylor’s law was observed, where the most expressed genes followed an exponential distribution, and the remaining genes followed a Poisson distribution. A biphasic behavior in scRNA-seq has been previously identified and were mainly attributed to the sampling process. For instance, shallow sequencing can mask the evidence of overdispersion which results in low expressing genes fitting to a Poisson distribution [20]. Interestingly, Lazzardi *et al*. found a triphasic Zipf’s law behavior in scRNA-seq data [4].

Overall, the infection course evolution of Taylor’s parameters between infected cells and the control simulated dataset was similar for hBECs, although infected hIECs present a distinct increase-decrease-increase behavior in comparison to the simulated datasets. This suggests that the progression of infection had a significant impact on the system dynamics of hIECs cells, whereas for hBECs, Taylor’s parameters were mostly influenced by sampling noise. Taylor’s parameter *V* has been used as a proxy to system stability in data from the human gut microbiota [12]. If *β* is constant across different samples, then changes in *V* correspond to variations to the standard deviation of all elements of the systems equally. If all elements display large standard deviation, we can assume that their rank is unstable. Here, however, both parameters *V* and *β* varied simultaneously along the course of infection, which might compromise the relationship between *V* and system stability. Nevertheless, our results show that infection with SARS-CoV-2 has a systemic effect on the properties of the distribution of transcripts at the cell level.

Whether there is a direct or indirect relationship between infection progression and Taylor’s parameters is inconclusive. One possibility is that in the absence of dropout (*i.e.*, all transcripts in a cell is sequenced), the whole system will better fit to an exponential distribution. In this case, it is likely that the observed change in the breakpoint as infection progresses is due to increase in noise in the system (here, not technical/sampling noise), meaning that the relationship is indirect. Another possibility is that the Poisson to exponential transition dynamics might arise from the interplay between RNA transcription bursts and RNA degradation, or as previously suggested, a suppression in the export of newly transcribed RNA out of the nucleus that will be latter degraded [3], which is affected by viral infection. The *RSI* of most genes was low, which is consistent with constant rank hopping along the course of infection due to transcriptional bursts and high rates of RNA degradation. The most expressed genes, however, displayed high *RSI* values, suggesting higher stability and lower RNA degradation rates. These genes followed the exponential distribution, which can be interpreted as aggregation behavior (Fig 1). Nevertheless, our results suggest that, at least for hIECs cells, the switch from a Poisson to exponential distribution and Taylor’s parameters are not only influenced by sampling noise but also by the progression of the disease, revealing that the whole system dynamics of transcripts at the cellular level is affected, directly or indirectly, by viral infection.

Several ribosomal proteins and some genes related to cellular respiration, protein folding, apoptosis and immune response showed signatures of punctual rank stability and/or persistent behavior. Mitochondria-encoded genes are likely not responding to infection given that, along the progression of infection, they always followed an exponential distribution even when their expression decreased (Fig 2B). However, some nuclear genes related to cellular respiration indeed showed signals of punctual stability and/or persistent behavior in all cell types. The protein product of ORF9b of both SARS-CoV-1 and -2 localizes to the mitochondria and interacts with the translocase of outer membrane (TOM) protein 70 (TOM70), a receptor involved in mitochondrial antiviral signaling and apoptosis [22], to suppress the cellular immune defense [23]. In line with this, the chaperone HSP90AA1, that interacts with TOM70 to induce apoptosis [23], showed signature of punctual stability in all cell types; while its paralog, HSP90AB1, showed signature of persistent rank behavior. Additionally, other proteins that are part of the TOM complex in the mitochondria, such as TOM5, TOM6, TOM7 and TOM20 displayed evidence of persistent rank behavior in all cell types.

Focusing on some genes that are known to be associated with COVID-19, we found that the C-X-C motif ligand chemokine genes *CXCL1* and *CXCL3*; the interferon stimulated gene *IFIT2*; the transcription factor *IRF1*; the AP-1 transcription factor proteins JUN and JUND; and the NF-κB inhibitor genes *NFKBIA*, *NFKBIZ* and *TNFAIP3*, showed evidence of punctual rank stability. Both *CXCL1* and *CXCL3* were found to be upregulated in response to SARS-CoV-2 infection [24]. *IFIT2* showed a bimodal expression pattern in immune cell types from patients with severe COVID-19 [25]. Interestingly, the bimodal expression of *IFIT2* should resemble aggregation behavior in these datasets. IRF1 regulates the expression of MHC class I, and was shown to be downregulated by SARS-CoV-2 ORF6-encoded protein [26]. JUN was found to be a hub in the SARS-CoV-2-host interactome [27], and both *JUN* and *JUND* showed signal of abnormal behavior in the same datasets used in the present study [3]. Lastly, the NF-κB signaling pathway is activated upon infection with SARS-CoV-2 and triggers inflammation and the production of cytokines [28,29]. Higher expression levels of *NFKBIA* and *TNFAIP3* in basal, ciliated and T cells were associated with the severity of COVID-19 [25]; and an insertion homozygosis of the *NFKBIZ* gene is associated with higher mortality by COVID-19 [30]. It is important to note, however, that in the datasets used in our study, it is likely that some infected cells were already responding to interferon and other immune signaling proteins from other previously infected cells. Thus, the punctual stability of some genes related to immune response may not be due to the cellular infection itself, but rather due to response to other infected cells. Additionally, given the higher-than-expected number of infected cells detected here, it is likely that some uninfected bystander cells are present at the beginning of the infection, meaning that the infection progression analyzed here starts at a point prior to infection.

Abnormal dynamics of ribosomal proteins and a few genes related to immune response in SARS-CoV-2-infected cells was previously detected in the same datasets used in this study [3]. Recently, an inverse relationship between inflammation and ribosome level was found, and furthermore, an increase in inflammation and decrease in ribosome level was associated with the severity of COVID-19 symptoms [31]. However, it remains unclear whether the ribosome content-inflammation interplay along the course of cellular infection bears any relevance to the dysregulated immune response associated with COVID-19 severity. Nevertheless, ribosomal proteins are tantalizing therapeutic targets due to their importance to viral translation as it has been recently shown that two ribosome inactivating proteins can inhibit SARS-CoV-2 replication in human lung epithelial cells (A549) [32]. In addition to ribosomal proteins, some translation initiation factors also showed evidence of punctual stability and/or persistent behavior. EIF3A and EIF3F, which are involved in the IRES-dependent translation of hepatitis C virus [33,34], showed, respectively, signature of persistent rank behavior and evidence of punctual stability and strong persistent rank and expression behavior; *EIF3E* showed evidence of punctual stability and persistent rank behavior; and several other translation initiation factors, namely *EIF1*, *EIF2AK2*, *EIF3K*, *EIF4G2*, and *EIF5*, showed evidence of persistent rank behavior. The RNA-binding activity of several components of *EIF3* is inhibited by SARS-CoV-2, which is in agreement with the role of SARS-CoV-2 NSP1 in inhibiting the recruitment of 40S to cellular mRNAs [35].

## Conclusion

Here, we successfully applied statistical frameworks from complex systems to scRNA-seq data to investigate the dynamics of cells infected with SARS-CoV-2 at the system and individual gene levels. Our results suggest a cell type-dependent systemic instability in response to SARS-CoV-2 infection. In hIECs, SARS-CoV-2 infection led to an increase, decrease and final increase in system stability (Fig 2A). In contrast, for hBECs, infection and sampling noise seemingly had the same effect on systemic instability (Fig 2A). Despite this systemic cell type-dependent response, several genes involved in translation, cellular respiration, apoptosis, protein-folding, and immune response showed evidence of deterministic behavior in all three cell types along the course of infection in the form of punctual rank stability or persistent rank behavior.

## Methods

### Data collection

Processed scRNA-seq data of human bronchial epithelial cells (hBECs) [18] and human intestinal epithelial cells (hIECs) from colon and ileum intestinal organoids [19] were obtained from [3]. The obtained processed gene frequencies matrices were previously generated by transforming UMI counts to transcript abundances. Briefly, UMI counts were modeled under a Poisson distribution, where transcript abundances were represented as the weighted average of transcript frequencies based on a normalized likelihood function [3]. Cells with at least 10 uncorrected viral UMIs were considered to be infected, and cells from mock data were considered to be uninfected. Infected cells were ordered based on their percentage of viral RNA, which is used here as a proxy of infection time and thus provide a measure of pseudotime of infection progression. Viral RNA counts were removed from the count matrices before downstream analyses, meaning that gene abundances were calculated using only cellular transcripts.

### Fit to Taylor’s law

To analyze the progression of infection through infection, cells were first ordered based on the accumulation of viral RNA then separated in 30 bins containing a similar number of cells (∼105, ∼147 and ∼124 cells for hBECs, colon and ileum cells, respectively) with a similar viral load. Genes exhibiting more than 95% of zeros were filtered out. The mean expression and standard deviation of each gene were calculated over the 30-bins based on their abundances in each cell. Then, Taylor’s parameters were estimated by fitting the log of means and standard deviations to a linear regression. The segmented R package v1.6-4 [36] was used to fit the log-transformed data to a segmented linear regression with one breakpoint. When fitting binned data to a segmented regression, mitochondrial genes and some selected nuclear genes were removed given that they always fit to an exponential distribution regardless of their mean expression, and therefore, they are likely not responding to infection and could influence the estimation of the parameters at some specific bins. Additionally, for binned data only, genes with more than 70% of zeros were filtered out when fitting the data to a biphasic model with one breakpoint. A simple schematic of the structure of the data used to estimate Taylor’s parameters is shown in S3A Fig, and S3B Fig shows a representation of the binned data used to investigate the progression of Taylor’s parameters along the course of infection.

### Simulation of increasing technical noise in uninfected cells

A down-sampled dataset was created for each cell type to simulate the expected increase in cellular transcript dropout due to viral RNA accumulation. To create a simulated cell, the transcriptional profile of an uninfected cell was used to randomly sample *n* transcripts, where *n* corresponds to the total number of cellular UMIs from an infected cell, and the probability of sampling a transcript from a given gene is its abundance in the uninfected cell. Sampling transcripts based on the gene abundances of an uninfected cell and the number of cellular UMIs from an infected one will create a simulated cell that will inherit the transcriptional profile of the uninfected cell and the sampling noise of the infected cell. Simulated cells are ordered based on the viral RNA accumulation of the infected cells that were used to simulate their sampling noise, and for each cell type, there are as much simulated cells as there are infected cells. The Seurat package v4.3.0 [37] was used for downstream analyses of simulated and uninfected cells. Counts were log-normalized, and a standard clustering analysis was performed, where the top ten principal components (PCs) were used for clustering and uniform manifold approximation and projection (UMAP) dimensional reduction. When dividing the simulated data into 30 bins, genes with more than 95% of zeros were filtered out before fits to Taylor’s law. For this simulated data, the progression through pseudotime should reflect increase in sampling noise.

### Gene rank stability

The rank stability index (*RSI*) of one gene is defined from its rank, determined from abundances matrices of cells ordered according to their viral load. Due to the high prevalence of zeros due to dropout, ties were resolved by randomization to avoid overestimation of stability of the less expressed genes. The *RSI* of each gene was computed based on its observed rank hops, *D*, (*i.e.*, the sum of the absolute number of rank differences between ordered adjacent cells) divided by the number of total possible rank hops as *RSI* = 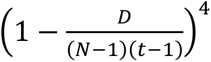, where *N* is the number of genes (rows), *t* is the number of cells (columns), and the power index is arbitrarily chosen to increase the resolution of stable elements [12]. S3C Fig shows a representation of a matrix containing the rank of each gene that was used to calculate their associated *RSI*.

### Estimation of punctual rank stability

To investigate signals of punctual rank stability, *RSI* values were calculated from the mean gene abundance at each bin instead of individual cells. Only genes that were expressed in at least one cell in every bin were further analyzed. Genes that displayed punctual stability, *i.e*., that presented higher stability at some point along the infection, were determined based on a resampling strategy with 1000 replicates. In addition, for each replicate, an *RSI* was calculated from a matrix where the order of the bins was shuffled, with the exception of the first and last ones. The probability of finding an *RSI* value at least as high as the observed *RSI* of a given gene was calculated by applying a survival function (1 – empirical cumulative distribution function) estimated from the *RSI* values calculated from shuffling. Genes with *RSI* values with a false discovery rate (FDR) < 0.05 were considered to have undergone through a change in their rank stability at some point throughout the course of infection. A punctual stability index (*PSI*) was calculated by dividing the gene *RSI* by the mean *RSI* of the replicates, where *PSI* > 1 is indicative of punctual stability. A schematic representation of the data transformation that was employed to estimate the punctual rank stability of each gene is shown in S3D Fig.

### Persistent behavior of gene rank

Long-range dependence and persistent behavior along the course of infection was investigated by estimating the Hurst exponent *H* for each gene separately for each cell type. A detailed explanation of the rescaled range analysis is available in S1 Appendix. Here, gene rank (see *Gene rank stability*; S3C Fig) was used to estimate *H* with the R package pracma v2.4.2 [38]. The robustness of this analysis was assessed by also estimating *H* for a set of control datasets that included the simulated datasets, uninfected cells, infected cells where cells were shuffled (and thus not ordered according to viral RNA accumulation) and a random matrix with the same number of rows (genes) and columns (cells) as the infected matrix where each value was drawn from a uniform distribution within the range [–1, 1]. A minimum window size of 50 was used when estimating *H* using the gene rank data. Genes with persistent behavior that simultaneously showed an *H* ≥ 0.7 in the infected dataset and *H* < 0.7 in its respective simulated dataset were further investigated.

### Gene ontology (GO) analyses

All gene set enrichment analyses were performed with the R packages clusterProfiler v4.8.2 [39] and org.Hs.eg.db v3.17.0 [40].

## Acknowledgements

This work was supported by CSIC PTI Salud Global grant 202020E153, by grants SGL2021-03-009 and SGL2021-03-052 from European Union NextGenerationEU/PRTR through the CSIC Global Health Platform established by EU Council Regulation 2020/2094, and by grants PID2022-136912NB-I00 funded by MCIU/AEI/10.13039/501100011033 and by “ERDF a way of making Europe”, and CIPROM/2022/59 funded by Generalitat Valenciana to S.F.E. J.A.O. work was partially supported by grant PID2019-109592GB-I00 from MCIU/AEI/10.13039/501100011033 and “ERDF a way of making Europe” and by Generalitat Valenciana grant CIAICO/2021/180.

## Data availability

All the R code used to generate these results are available at https://github.com/jmfagundes/sarscov2scrna.

## Supporting information captions

**S1 Fig. Characteristics of simulated datasets.**

(a) For each cell type, boxplots of the proportion of zeros of each gene for each bin for the infected and simulated dataset. (b) UMAP projections of the infected and simulated datasets. Each point represents a cell.

**S2 Fig. Taylor’s parameters of the biphasic fit for binned data per cell.**

**S3 Fig. Schematic representation of the data structure used in each analysis.**

(a) Representation of a gene abundance matrix (left) that was log-transformed and fitted to a linear regression (right) to estimate Taylor’s parameters. (b) Pseudotime binned data (left). Infected cells were sorted into bins so that the viral load of any cell in bin *i* is lower than the viral load of any cell in bin *i* + 1. Taylor’s parameters were estimated for each bin (right). (c) Representation of a gene rank matrix used for the calculation of *RSI* shown in Fig 1 and for the estimation of the Hurst exponent of each gene. (d) Mean gene abundances of each bin (left) were used to generate a rank matrix (right) from which punctual rank stability analyses were conducted.

**S1 File. Results of the rank stability dynamics analyses.**

The *RSI,* mean *RSI* of a 1000 replicates, *P*-value and adjusted *P*-value (FDR) and *PSI* of each gene for each dataset is shown in separate sheets. The last sheet corresponds to the GO enrichment analysis of the genes that exhibited signal of punctual rank stability concomitantly in all three cell types.

**S2 File. Results of the R/S analyses.**

The empirical *H* exponent of each gene for each dataset is shown in separate sheets for each cell type. The last sheet corresponds to the GO enrichment analysis of the genes that exhibited signal of strong persistent behavior concomitantly in all three cell types.

**S1 Appendix. Detailed explanation of R/S analysis.**

