## Appendix S1 for "System and transcript dynamics of cells infected with severe acute respiratory syndrome virus 2 (SARS-CoV-2)"

### Appendix: Rescaled Range Analysis

Let $g_i$, with $i = 1, 2, \ldots, n$, denote a normalized sequence of genetic expressions along the time evolution. The index $i$ stands for pseudo-time. We define the mean-adjusted sequence $h_i$, with $i = 1, \ldots, n$, as

$$h_i^{(n)} = g_i - G_n, \quad \text{with} \quad G_n = \frac{1}{n}\sum_{k=1}^{n} g_k. \tag{1}$$

We will consider sub-sequences of length $\tau < n$ and denote every single one as $h_i^{(\tau)}$, with the index $i$ shifted such that $1 \leq i \leq \tau$. In addition, we compute the cumulative deviates series

$$z_j^{(\tau)} = \sum_{k=1}^{j} h_k^{(\tau)}. \tag{2}$$

Eventually, the range $R(\tau)$ of a sub-sequence is obtained as

$$R(\tau) = \max_{1\leq i\leq \tau}\{h_i^{(\tau)}\} - \min_{1\leq i\leq \tau}\{h_i^{(\tau)}\}, \tag{3}$$

and the associated standard deviation of the $\tau$ observations in the window reads

$$S(\tau) = \left[\frac{1}{\tau}\sum_{k=1}^{\tau}\left(h_k^{(\tau)}\right)^2\right]^{1/2}, \tag{4}$$

Hurst observed empirically that for a number of real data records the rescaled range has a power law behavior

$$R(\tau)/\langle S(\tau) = (\tau/\alpha)^H, \quad \alpha > 0. \tag{5}$$

The important point is that the Hurst exponent, $0 < H < 1$, admits an interesting interpretation. For records generated by statistically independent processes with finite variance the rescaled range behaves asymptotically as

$$R(\tau)/S(\tau) = (\pi\tau/2)^{1/2}. \tag{6}$$

Instead, sequences that give rise to an $R/S$ power law with an exponent $H \neq 1/2$ point out the presence of long range correlation in the series. The case $1/2 < H$ is said persistent and means that large (irrespective, small) values are more likely followed by large (irrespective, small) values in the sequence. The case $H < 1/2$ is termed antipersistent and presents the opposite behavior.

It turns out that the Hurst empirical law (5), introduced in a context of hydrology, gives a value close to $H = 0.73$ in a number of phenomena in Nature recorded as numerical series. This is referred to as *Hurst phenomenon* in the literature [1].

A simple way to estimate $H$ from data is as the slope of the linear regression of $\log[R(\tau)/S(\tau)]$ vs. the window size $\log\tau$: $\log(R/S) = H\,\log\tau - H\log\alpha$, as in Figure 1.

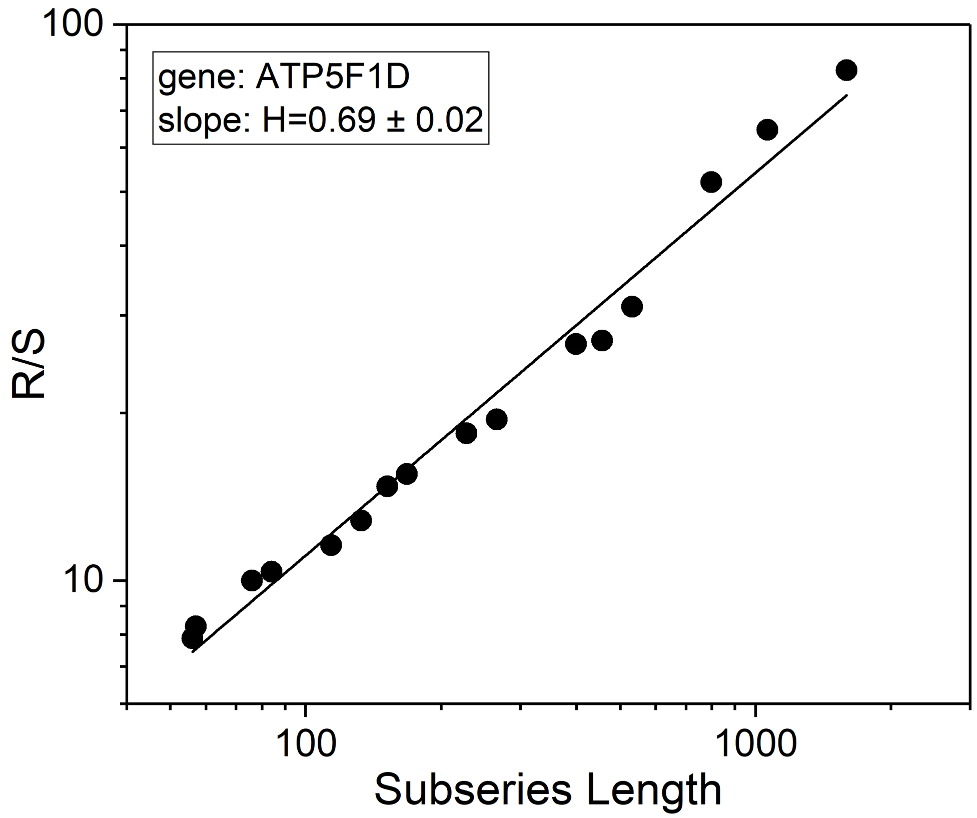


Figure 1: $R/S$ analysis of gene ATP5F1D from normalized expression. The Hurst exponent $H$ is the slope of the best fit line in log-log scale.
