## Supplementary figures and images for "System and transcript dynamics of cells infected with severe acute respiratory syndrome virus 2 (SARS-CoV-2)"

### Supplementary Fig. S1

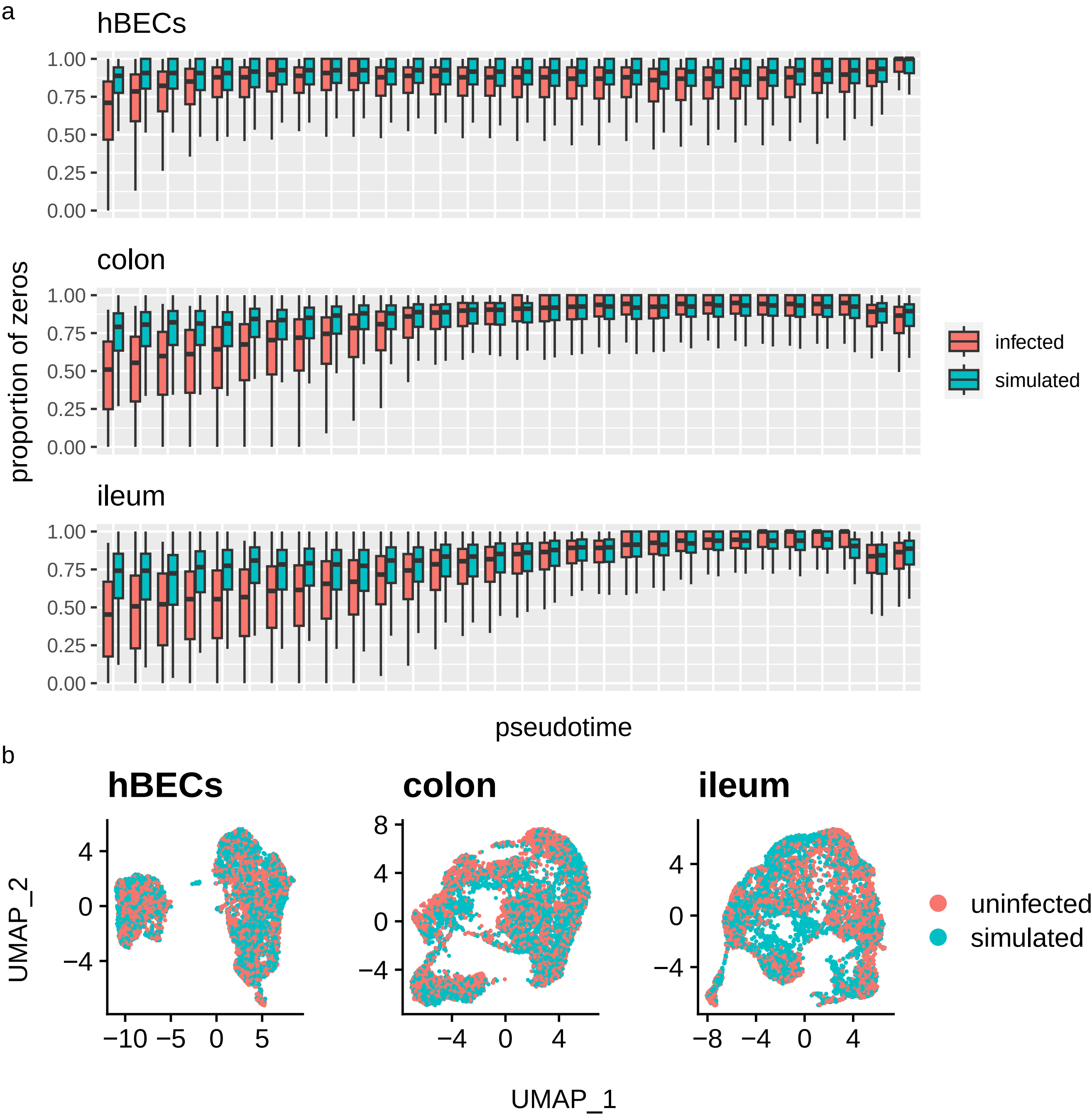

### Supplementary Fig. S2

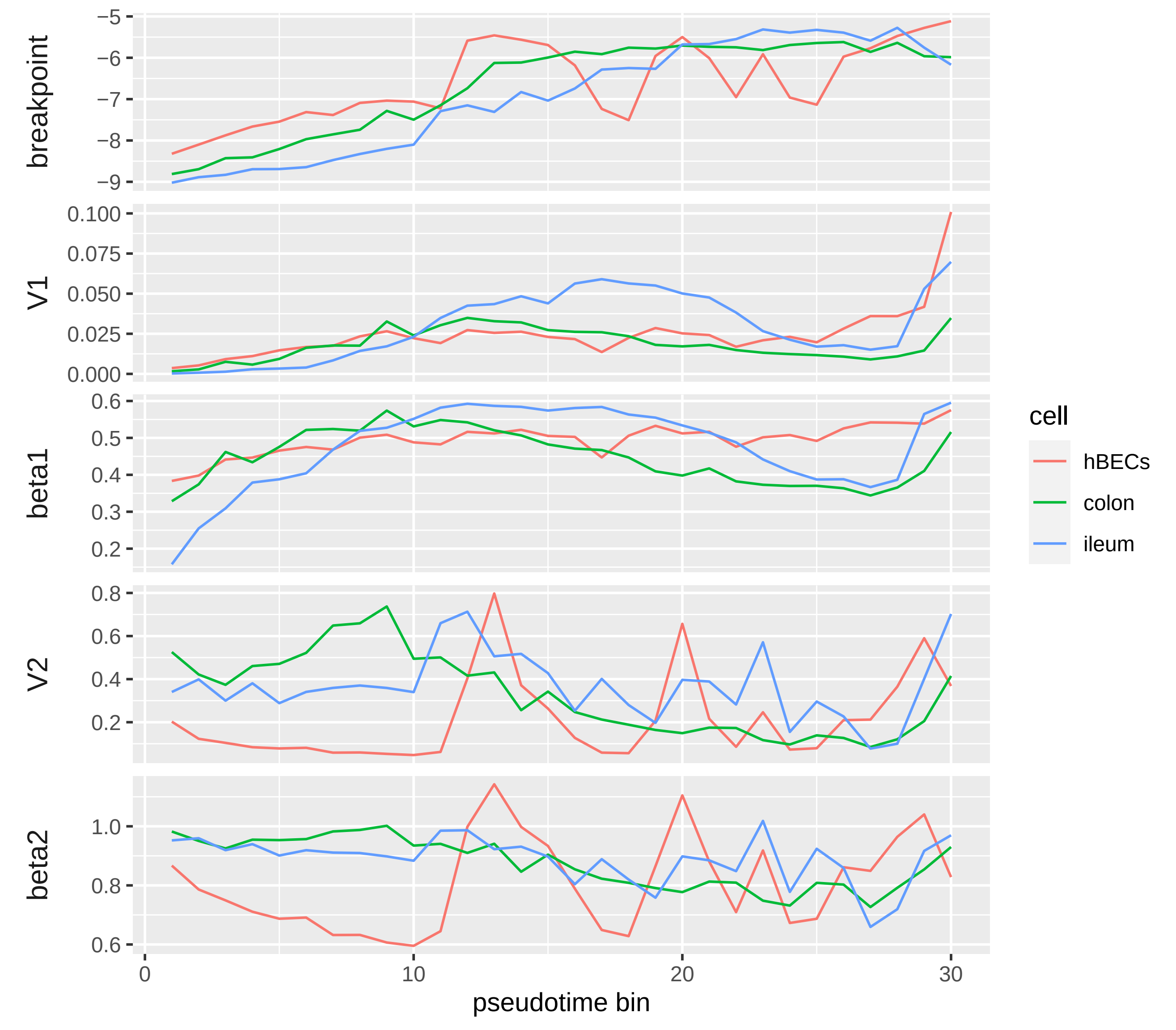
