## Supplementary Fig. S3 for "System and transcript dynamics of cells infected with severe acute respiratory syndrome virus 2 (SARS-CoV-2)"

a

| | cell i | cell i + 1 | ... |
|---|---|---|---|
| *gene a* | 0.00011 | 0.00039 | |
| *gene b* | 0.00083 | 0.00121 | |
| *gene c* | 9e−05 | 0 | |
| ... | | | |

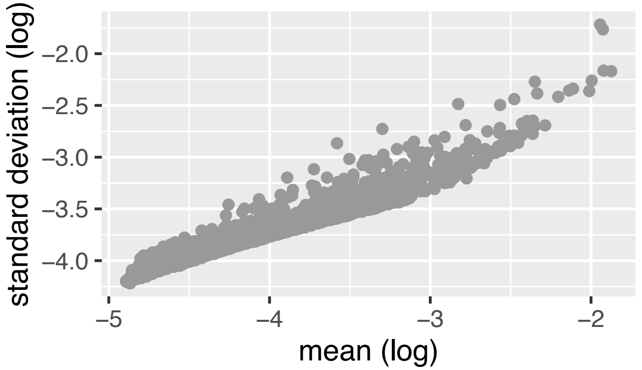


b

bin i

| | cell 1 | ... | cell 100 |
|---|---|---|---|
| *gene a* | 0.00011 | | 0.00032 |
| *gene b* | 0.00083 | | 0.001 |
| *gene c* | 9e−05 | | 0 |
| ... | | | |

bin i + 1

| | cell 101 | ... | cell 200 |
|---|---|---|---|
| *gene a* | 2e−04 | | 0.00011 |
| *gene b* | 0.0011 | | 0.00124 |
| *gene c* | 0 | | 0 |
| ... | | | |

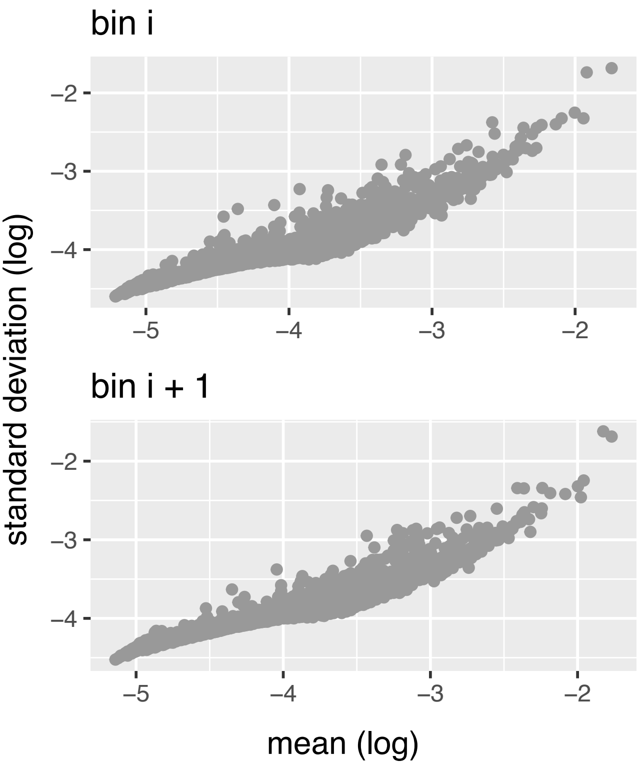


c

| | cell i | cell i + 1 | ... |
|---|---|---|---|
| *gene a* | 1518 | 290 | |
| *gene b* | 160 | 114 | |
| *gene c* | 3520 | 6153 | |
| ... | | | |

d

mean gene abundances

| | cells 1−100 | cells 101−200 | ... |
|---|---|---|---|
| *gene a* | 0.00026 | 0.00029 | |
| *gene b* | 0.00096 | 0.00103 | |
| *gene c* | 1e−05 | 1e−05 | |
| ... | | | |

rank of mean gene abundances

| | cells 1−100 | cells 101−200 | ... |
|---|---|---|---|
| *gene a* | 362 | 788 | |
| *gene b* | 118 | 110 | |
| *gene c* | 4304 | 5776 | |
| ... | | | |
